# S protein-reactive IgG and memory B cell production after human SARS-CoV-2 infection includes broad reactivity to the S2 subunit

**DOI:** 10.1101/2020.07.20.213298

**Authors:** Phuong Nguyen-Contant, A. Karim Embong, Preshetha Kanagaiah, Francisco A. Chaves, Hongmei Yang, Angela R. Branche, David J. Topham, Mark Y. Sangster

**Affiliations:** David H. Smith Center for Vaccine Biology and Immunology, Department of Microbiology and Immunology, University of Rochester Medical Center, Rochester, New York, USA; Department of Biostatistics and Computational Biology, University of Rochester Medical Center, Rochester, New York, USA; Department of Medicine, University of Rochester Medical Center, Rochester, New York, USA

**Keywords:** SARS-CoV-2, COVID-19, Memory B cells, IgG antibodies, Spike protein

## Abstract

The high susceptibility of humans to SARS-CoV-2 infection, the cause of COVID-19, reflects the novelty of the virus and limited preexisting B cell immunity. IgG against the SARS-CoV-2 spike (S) protein, which carries the novel receptor binding domain (RBD), is absent or at low levels in unexposed individuals. To better understand the B cell response to SARS-CoV-2 infection, we asked whether virus-reactive memory B cells (MBCs) were present in unexposed subjects and whether MBC generation accompanied virus-specific IgG production in infected subjects. We analyzed sera and PBMCs from non-SARS-CoV-2-exposed healthy donors and COVID-19 convalescent subjects. Serum IgG levels specific for SARS-CoV-2 proteins (S, including the RBD and S2 subunit, and nucleocapsid [N]) and non-SARS-CoV-2 proteins were related to measurements of circulating IgG MBCs. Anti-RBD IgG was absent in unexposed subjects. Most unexposed subjects had anti-S2 IgG and a minority had anti-N IgG, but IgG MBCs with these specificities were not detected, perhaps reflecting low frequencies. Convalescent subjects had high levels of IgG against the RBD, S2, and N, together with large populations of RBD- and S2-reactive IgG MBCs. Notably, IgG titers against the S protein of the human coronavirus OC43 in convalescent subjects were higher than in unexposed subjects and correlated strongly with anti-S2 titers. Our findings indicate cross-reactive B cell responses against the S2 subunit that might enhance broad coronavirus protection. Importantly, our demonstration of MBC induction by SARS-CoV-2 infection suggests that a durable form of B cell immunity is maintained even if circulating antibody levels wane.

**IMPORTANCE:** Recent rapid worldwide spread of SARS-CoV-2 has established a pandemic of potentially serious disease in the highly susceptible human population. Key questions are whether humans have preexisting immune memory that provides some protection against SARS-CoV-2 and whether SARS-CoV-2 infection generates lasting immune protection against reinfection. Our analysis focused on pre- and post-infection IgG and IgG memory B cells (MBCs) reactive to SARS-CoV-2 proteins. Most importantly, we demonstrate that infection generates both IgG and IgG MBCs against the novel receptor binding domain and the conserved S2 subunit of the SARS-CoV-2 spike protein. Thus, even if antibody levels wane, long-lived MBCs remain to mediate rapid antibody production. Our study also suggests that SARS-CoV-2 infection strengthens preexisting broad coronavirus protection through S2-reactive antibody and MBC formation.

## INTRODUCTION

The betacoronavirus SARS-CoV-2, the causative agent of a respiratory disease termed COVID-19, emerged in China in late 2019 and rapidly spread worldwide (1). A pandemic was declared in March 2020 and global deaths from COVID-19 now exceed 500,000. The rapid increase in cases in many countries has challenged healthcare systems and shutdowns and quarantine measures introduced to slow virus spread have caused major disruptions to society and economies (2). SARS-CoV-2 infection produces a wide spectrum of outcomes. A proportion of infections, likely more than 20%, remain asymptomatic. Most clinical cases develop mild to moderate respiratory symptoms, but up to 20% progress to a more severe disease with extensive pneumonia (3, 4).

When SARS-CoV-2 emerged and began to spread, the severity of the threat was primarily attributed to the novelty of the virus to the human immune system and, consequently, a lack of preexisting immune memory to quickly clear virus and limit disease progression. Four types of common cold coronavirus are endemic in humans, the alphacoronaviruses 229E and NL63 and the betacoronaviruses OC43 and HKU1. However, limited relatedness between key structural proteins of these human coronaviruses (HCoVs) and those of SARS-CoV-2 suggested that significant cross-reactive immunity was unlikely (5, 6). Initial studies of non-SARS-CoV-2-exposed individuals found negligible levels of IgG against the SARS-CoV-2 spike (S) protein, the viral attachment protein that binds the receptor angiotensin converting enzyme 2 (ACE2) on host cells to initiate infection (7). More recently, however, studies have provided evidence of SARS-CoV-2-reactive B and T cell memory in unexposed subjects that could confer some protection against SARS-CoV-2 or modulate disease pathogenesis.

Sera from non-SARS-CoV-2-exposed individuals have been screened for IgG binding to the S1 and S2 subunits of the SARS-CoV-2 S protein. The membrane-distal S1 subunit contains the receptor binding domain (RBD) for receptor recognition, and the membrane-proximal S2, which has higher homology among coronaviruses than does S1 (6, 8), mediates membrane fusion to release viral RNA into the host cell. In two large cohorts of unexposed subjects, approximately 10% had IgG that bound S2, but not S1 or the RBD. Approximately 4% of subjects had IgG against the SARS-CoV-2 nucleocapsid (N) protein, which is highly conserved among coronaviruses (9, 10). Although N is an internal viral protein and not a target of neutralizing antibodies (Abs), coronavirus infections typically elicit strong anti-N Ab production (11). The idea that circulating HCoVs elicit IgG that cross-reacts with SARS-CoV-2 is supported by the finding that SARS-CoV-2 infection increases IgG titers against the S proteins of multiple HCoVs (12). In T cell studies, CD4^+^ T cells in up to 50% of non-SARS-CoV-2-exposed donors responded to epitopes in S and non-S proteins of SARS-CoV-2 (8, 13). Notably, S-reactive CD4^+^ T cells in unexposed subjects were mostly reactive to the conserved S2 subunit, consistent with cross-reactivity to circulating HCoVs (8). SARS-CoV-2-reactive CD8^+^ T cells were also detected in unexposed donors, but the response was less marked than for CD4^+^ T cells (13).

SARS-CoV-2-reactive memory B cells (MBCs) generated in B cell responses to HCoVs are also likely to be present in non-SARS-CoV-2-exposed individuals. Indeed, MBCs might be more important than preexisting cross-reactive Abs as a source of protection against SARS-CoV-2. IgG MBCs are more broadly reactive than Abs generated against the same antigen, they persist after circulating Ab levels wane, and they are readily activated to generate strong Ab responses or seed germinal centers for additional rounds of affinity maturation (14). Concurrent early production of virus-specific IgM and IgG in the response to SARS-CoV-2 infection suggests a response mediated by IgG MBCs as well as naïve B cells (9, 15–17). This picture is supported by identification of B cell subsets with high and low immunoglobulin V gene mutation frequencies during the response to SARS-CoV-2 infection (18). However, little direct analysis of SARS-CoV-2-reactive MBCs in unexposed subjects has been performed.

Characterization of MBCs generated and/or expanded by SARS-CoV-2 infection can also provide insights into cross-reactivity between coronaviruses and participation of preexisting MBCs in the response. Wec et al. (19) used cells from a survivor of the 2003 SARS-CoV outbreak as a source of MBCs that bound the S protein of SARS-CoV-2; a comprehensive panel of Abs expressed by the MBCs were cloned and characterized. Notably, most of the highly mutated mAbs bound the S2 subunit of multiple HCoV S proteins, often with higher affinity than to the S2 of SARS-CoV-2. A screening of healthy donors identified low frequencies of MBCs reactive to the S proteins of the 2003 SARS-CoV and SARS-CoV-2 (19). Findings suggest that S2-reactive MBCs generated by HCoVs were activated and expanded by the 2003 SARS-CoV. RBD-binding MBCs sampled in the convalescent phase of SARS-CoV-2 infection expressed Abs with relatively low numbers of V gene mutations, suggesting that this component of the response largely reflected naïve B cell activation by novel epitopes (20).

To extend our understanding of the B cell response to SARS-CoV-2 infection, the current study compared Ab and MBC immunity to SARS-CoV-2 in unexposed individuals and individuals in the convalescent phase of infection. In particular, we were interested in the presence of SARS-CoV-2-reactive MBCs in unexposed subjects that could confer some protection against SARS-CoV-2, and formation of MBCs by SARS-CoV-2 infection to provide durable protection against reinfection. Most importantly, we demonstrate that SARS-CoV-2 infection generates both IgG and IgG MBCs reactive to the novel RBD and the conserved S2 subunit of the S protein. Long-lived MBCs are thus likely to be available to mediate rapid protective Ab responses if circulating Ab levels wane and reinfection occurs. Our study also draws attention to preexisting SARS-CoV-2-cross-reactive B cell memory to the S2 subunit in SARS-CoV-2-naïve subjects. We speculate that the strong response to S2 after SARS-CoV-2 infection reflects preexisting S2-reactive MBC activation and strengthens broad coronavirus protection.

## RESULTS

### IgG against SARS-CoV-2 proteins in unexposed subjects primarily targets the S2 subunit of the S protein

To investigate preexisting B cell immunity to SARS-CoV-2 in unexposed individuals and SARS-CoV-2-reactive B cell immunity generated by infection, we analyzed sera and peripheral blood mononuclear cells (PBMCs) from (i) 21 healthy donors sampled prior to the emergence of SARS-CoV-2, and (ii) 26 non-hospitalized COVID-19 convalescent subjects sampled 4-9 weeks after symptom onset. Reactivity was measured against the S (including the RBD and S2 subunit) and N proteins of SARS-CoV-2 and the S proteins of the human alphacoronavirus 229E and betacoronavirus OC43. The H1 influenza virus hemagglutinin and tetanus toxoid (TTd) were included as control antigens that humans are commonly exposed to through infection and vaccination.

Serum IgG levels were measured by ELISA. Approximately one-third of non-SARS-CoV-2-exposed subjects in the healthy donor cohort had low levels of serum IgG against the S and N proteins of SARS-CoV-2, likely reflecting cross-reactivity with seasonal HCoVs (Figure 1A). Notably, 86% of unexposed subjects had IgG against the highly conserved S2 subunit of the S protein. It is possible that inherent features of the bulky S reagent used in our analysis reduced binding by anti-S2 Abs. IgG that bound the highly novel RBD was not detected in unexposed subjects. All non-SARS-CoV-2-exposed subjects had IgG against S proteins of the HCoVs 229E and OC43, indicating previous infection, and against the control proteins H1 and TTd (Figures 1C-1F).

**Figure 1.**
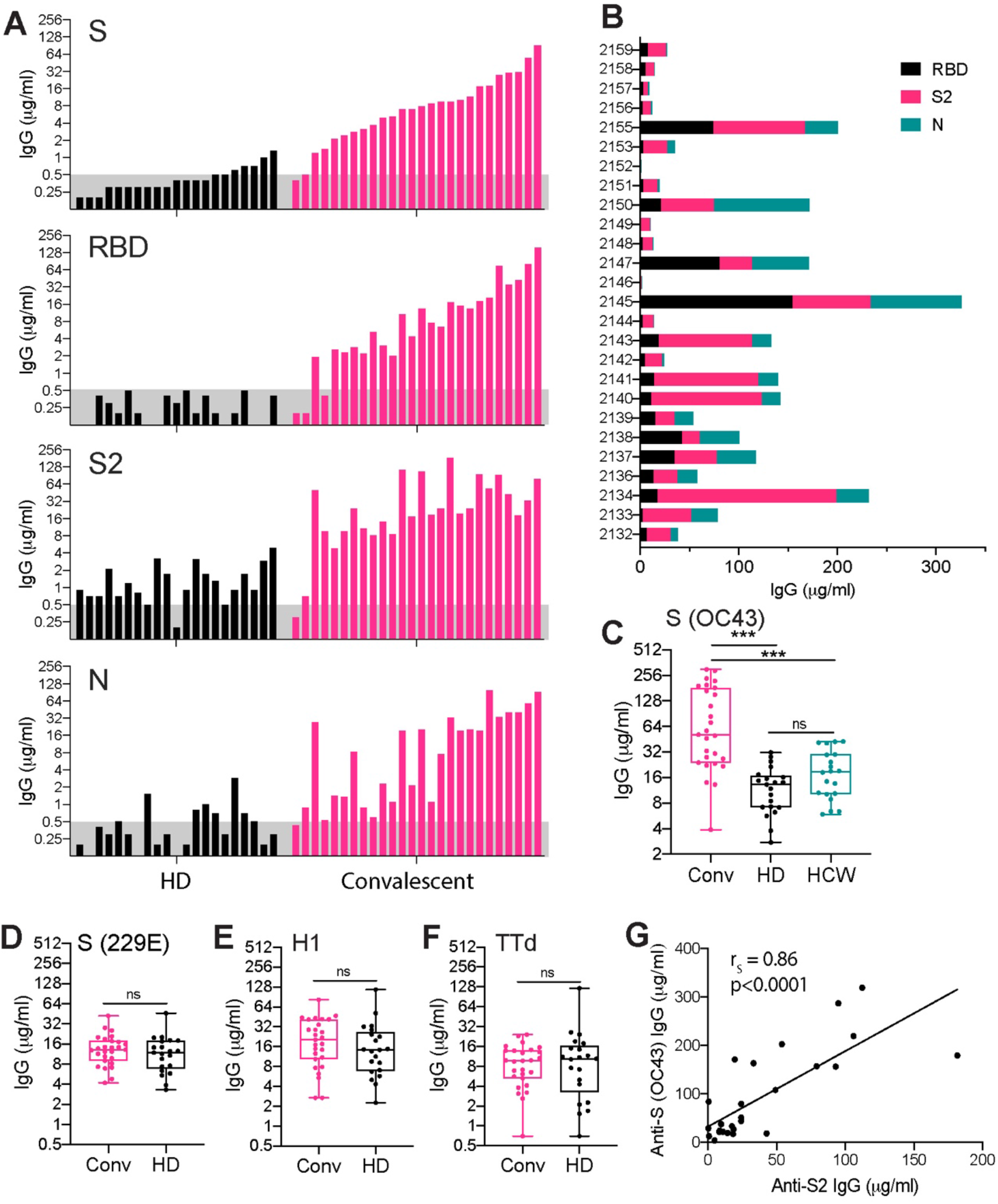
Serum IgG levels against SARS-CoV-2 and non-SARS-CoV-2 proteins in non-SARS-CoV-2-exposed and COVID-19 convalescent subjects. Sera were collected from (i) 21 healthy donors sampled from 2011-14 (HD), (ii) 20 SARS-CoV-2-negative healthcare workers sampled in 2020 (HCW), and (iii) 26 COVID-19 convalescent subjects sampled 4-9 weeks after symptom onset (CONV). (A) Serum IgG concentrations measured by ELISA against the SARS-CoV-2 spike (S), receptor binding domain (RBD), S2 subunit, and nucleocapsid (N). Columns represent individual HD and CONV subjects in order of ascending titers against S. The assigned cutoff for positivity is shown by the shaded bar. (B) Proportions of serum IgG against the SARS-CoV-2 RBD, S2, and N for individual CONV subjects. (C) Serum IgG concentrations against the S protein of the HCoV OC43 in CONV, HD, and HCW subjects. (D-F) Serum IgG concentrations against the S protein of the HCoV 229E (D), the influenza virus H1 hemagglutinin (E), and TTd (F) in CONV and HD subjects. (G) Correlation between serum IgG concentrations against the S2 subunit of SARS-CoV-2 and the S protein of the HCoV OC43. Significance (*, P < 0.05; **, P < 0.01; ***, P < 0.001; ns [not significant]) for comparisons of serum IgG concentrations between subject groups was determined by the Wilcoxon rank-sum test. Correlations were tested by Spearman correlation analysis with corresponding robust regression models.

### S- and N-specific IgG production following SARS-CoV-2 infection includes a strong response to the S2 subunit

Levels of IgG against S, RBD, S2 and N were markedly higher in convalescent subjects than unexposed subjects, indicating strong induction of these Abs by SARS-CoV-2 infection (Figure 1A). In a small number of convalescent subjects, high anti-S IgG titers were associated with low levels of anti-N IgG. Indeed, more than 40% of convalescent subjects had anti-N IgG levels within the range in unexposed subjects, questioning the reliability of using anti-N IgG measurement to identify previous SARS-CoV-2 infection.

Notably, serum IgG titers against S2 were consistently higher than against the RBD in convalescent subjects, perhaps reflecting the novelty of the RBD and a response dependent on naive B cell activation (Figure 1B). Interestingly, titers of IgG were higher against the S protein of the HCoV OC43 in convalescent subjects than in unexposed subjects, but this was not the case for the S protein of HCoV 229E (or for the control proteins H1 and TTd) (Figures 1C-1F). The anti-OC43 S IgG titers correlated with those against the SARS-CoV-2 S (r_S_ = 0.49, *P* = 0.0109), RBD (r_S_ = 0.57, *P* =0.0025), and S2 (r_S_ = 0.86, *P*<0.0001), indicating a relationship with SARS-CoV-2 infection (Figure 1G). The particularly strong correlation between IgG titers against OC43 S and the SARS-CoV-2 S2 suggests a cross-reactive response to the S2 subunit.

Since the healthy donor samples in our analysis were collected 6-10 years before the emergence of SARS-CoV-2, we considered the possibility that a recently circulating HCoV could have been responsible for the higher anti-OC43 S IgG titers in the convalescent subjects. To exclude this possibility, we measured anti-OC43 S IgG titers in sera collected from 20 healthcare workers in 2020. The healthcare workers cared for hospitalized SARS-CoV-2 patients, but all were negative for IgG against SARS-CoV-2 S and RBD, consistent with the effectiveness of personal protective equipment and appropriate work practices. OC43 S-reactive IgG levels in healthcare worker sera were similar to those in non-SARS-CoV-2-exposed healthy donor sera and significantly lower than those in sera from convalescent subjects (Figure 1C). Taken together, our results indicate that SARS-CoV-2 infection generates a strong IgG response that cross-reacts with the S2 of human betacoronaviruses.

### Strong S-reactive MBC formation following SARS-CoV-2 infection includes reactivity to the RBD and S2 subunit

PBMCs from non-SARS-CoV-2-exposed subjects and convalescent subjects were analyzed for MBCs reactive to SARS-CoV-2 proteins. Circulating antigen-specific IgG MBC populations were measured by in vitro stimulation of MBCs to induce differentiation into Ab-secreting cells (ASCs). Post-stimulation antigen-specific measurement of MBC-derived ASCs (MASCs) by ELISpot assay or MBC-derived polyclonal Abs (MPAbs) by ELISA provided a measure of the precursor MBCs (30). Analysis of MASCs by ELISpot assay was performed against the SARS-CoV-2 S, RBD, and N proteins, and the influenza H1 and TTd. MPAbs were measured against antigens used in the ELISpot assay, as well as SARS-CoV-2 S2, and the S proteins of the HCoVs OC43 and 229E. Antigen-specific IgG MPAb concentrations correlated strongly with the frequency of IgG MASCs derived from stimulated MBCs (determined for SARS-CoV-2 S and RBD, influenza H1, and TTd, r_S_ = 0.89, 0.67, 0.83, and 0.95, respectively, *P* ≤ 0.0002), validating MPAb concentration as a measure of the size of specific MBC populations.

The presence of a low level of IgG against the SARS-CoV-2 S, RBD, and N proteins in a proportion of unexposed subjects suggested that IgG MBCs with the same specificity had also been formed. However, these MBCs were not detected, possibly because of very low frequencies in the circulation. In contrast, IgG MBCs reactive to the S proteins of the HCoVs OC43 and 229E and the control proteins H1 and TTd were detected in nearly 50% or more of non-SARS-CoV-2-exposed subjects, consistent with the higher levels of serum IgG against these antigens (Figure 2E-2H). As expected, SARS-CoV-2 RBD-reactive MBCs were not detected in unexposed subjects.

**Figure 2.**
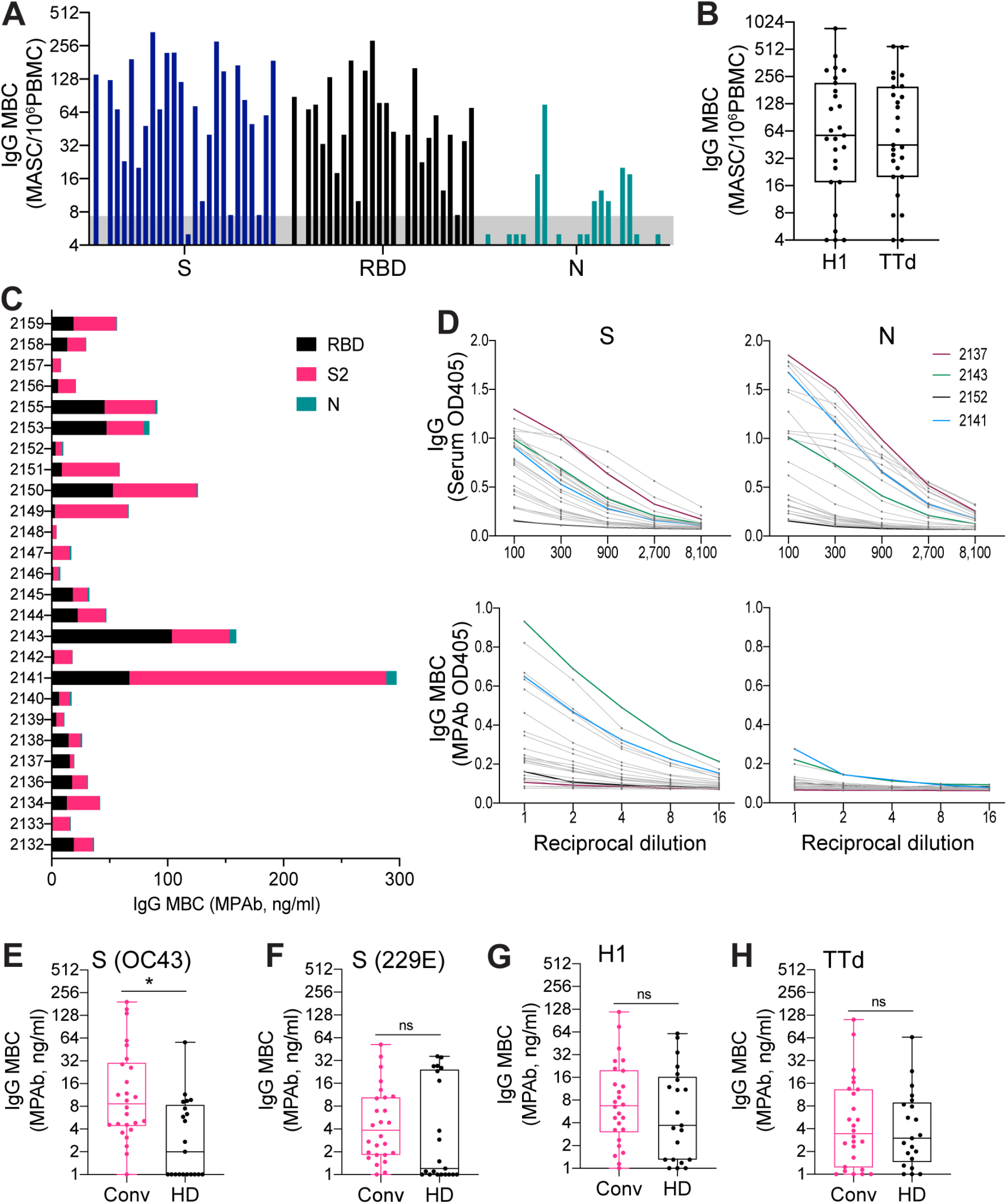
Analysis of IgG memory B cells (MBCs) reactive to SARS-CoV-2 and non-SARS-CoV-2 proteins in non-SARS-CoV-2-exposed and COVID-19 convalescent subjects. PBMCs for MBC analysis were collected from (i) 21 healthy donors sampled from 2011-14 (HD) and (ii) 26 COVID-19 convalescent subjects sampled 4-9 weeks after symptom onset (CONV). PBMCs were stimulated in vitro to induce MBC differentiation into Ab-secreting cells. Antigen-specific quantitation of MBC-derived Ab (IgG)-secreting cells (MASCs) or MBC-derived polyclonal (IgG) Abs (MPAbs) provided a measure of the abundance of specific IgG MBCs. (A) IgG MBCs reactive to the SARS-CoV-2 spike (S), receptor binding domain (RBD), and nucleocapsid (N) in CONV subjects. MBC numbers were determined by enumeration of IgG MASCs by ELISpot essay after in vitro MBC stimulation. The assigned cutoff for positivity is shown by the shaded bar. (B) IgG MBCs reactive to the influenza virus H1 hemagglutinin and TTd in CONV subjects. MBC numbers were determined by enumeration of IgG MASCs. (C) Proportions of IgG MBCs reactive to the SARS-CoV-2 RBD, S2, and N for individual CONV subjects. (D) Comparison of serum IgG concentrations (upper panels) and IgG MBC numbers (lower panels) reactive to the SARS-CoV-2 S (left-hand side) and N (right-hand side) proteins. Serum IgG was measured by ELISA; IgG MBC numbers were based on ELISA analysis of MPAbs. Dilution curves are shown for individual CONV subjects; curves for 4 subjects are shown in different colors to identify particular response patterns. (E-H) IgG MBCs reactive to the S proteins of HCoVs OC43 (E), and 229E (F), the H1 hemagglutinin (G), and TTd (H) in CONV and HD subjects. IgG MBC numbers were based on ELISA measurements of MPAbs. Significance (*, *P* < 0.05; ns [not significant]) for comparisons of IgG MBC numbers between subject groups was determined by the Wilcoxon rank-sum test.

In marked contrast to non-SARS-CoV-2-exposed subjects, the vast majority of convalescent subjects had circulating IgG MBCs reactive to the SARS-CoV-2 S, RBD, and S2, indicating strong induction by SARS-CoV-2 infection of MBCs reactive to novel and conserved regions of the S protein (Figure 2A). Notably, numbers of IgG MBCs reactive to the S protein of the HCoV OC43 were higher in convalescent subjects than in unexposed subjects (Figure 1E), but there was no difference between the two subject groups in IgG MBCs reactive to the HCoV 229E S protein, influenza H1 or TTd (Figures 2B, 2F-2H). S2-reactive IgG MBC numbers correlated well with IgG MBCs reactive to SARS-CoV-2 S (r_S_ = 0.77, *P* < 0.0001) and RBD (r_S_ = 0.60, *P* = 0.0012), and S of HCoV OC43 (r_S_ = 0.52, *P* = 0.0059), but not to S of HCoV 229E (r_S_ = −0.13, *P* = 0.53), influenza H1 (r_S_ = 0.13, *P* = 0.54), or TTd (r_S_ = 0.29, *P* = 0.15). The findings of our MBC analysis are consistent with serum IgG measurement and indicate that SARS-CoV-2 infection generates IgG MBCs reactive to the SARS-CoV-2 S2 that cross-react with the S2 of human betacoronaviruses. Interestingly, only a small proportion of the convalescent subjects generated detectable N-reactive IgG MBCs, even though most subjects produced high levels of anti-N IgG in serum (Figures 2C, 2D). It is unclear whether this reflects a real difference between S- and N-reactive MBC formation or an effect of the sampling time. Overall, we demonstrate that SARS-CoV-2 infection induces strong S-reactive MBC formation that would be expected to provide lasting protection against reinfection and potentially broad protection against betacoronaviruses.

## DISCUSSION

Our goals in this study were to investigate SARS-CoV-2-reactive B cell memory in unexposed subjects that could provide some protection against SARS-CoV-2 infection, and the generation of B cell memory by SARS-CoV-2 infection that could provide lasting protection against re-infection. In particular, we were interested in IgG MBCs, which respond to cognate antigens with rapid, vigorous, and high-affinity Ab production. Importantly, MBCs are long-lived cells that continue to provide strong protection when circulating Ab levels wane. Our approach was to analyze circulating IgG as well as IgG MBCs from the SARS-CoV-2-naïve and SARS-CoV-2-convalescent subject groups. Our key findings are as follows: (i) the presence of IgG reactive to the S2 subunit of SARS-CoV-2 in most unexposed subjects, likely reflecting cross-reactivity to HCoVs, (ii) markedly increased levels of IgG against the SARS-CoV-2 S and N proteins, including reactivity to the RBD and S2 subunit of S, in convalescent subjects, (iii) increased IgG binding to the S protein of the OC43 HCoV, but not 229E HCoV, in convalescent subjects, reflecting greater cross-reactivity between S2 subunits of betacoronaviruses, (iv) strong formation of IgG MBCs reactive with the RBD and S2 subunit of the SARS-CoV-2 S protein in convalescent subjects, and (v) formation of IgG MBCs reactive with the S protein of OC43, but not 229E, in convalescent subjects, consistent with S2 subunit cross-reactivity between betacoronaviruses.

Approximately one-third of our cohort of non-SARS-CoV-2-exposed subjects had low levels of IgG against the SARS-CoV-2 S and N proteins. The anti-N IgG likely reflects infection with HCoVs, which have low level (20-30%) homology with the SARS-CoV-2 N protein (10). However, a protective function for anti-N Abs has not been established (21). Notably, 86% of unexposed subjects had IgG against the S2 subunit, reflecting homology with HCoVs, but none had IgG against the highly novel SARS-CoV-2 RBD (6, 8, 22). Abs that target the S2 subunit have been shown to have virus neutralizing activity, raising the possibility that preexisting anti-S2 IgG confers some protection against SARS-CoV-2 (23). The processes that generate anti-S2 IgG are also likely to generate S2-reactive IgG MBCs and these might provide more significant protection than low levels of anti-S2 Abs. However, S2-reactive MBCs (or S-reactive and N-reactive MBCs) were not detected in non-SARS-CoV-2-exposed subjects. Taken together with the identification of S-reactive MBCs in unexposed healthy donors (19), it is likely that S2-reactive MBCs were below the limit of detection in our assays. Most MBCs are resident in lymphoid tissues, except for MBCs against frequently seen immunogenic antigens (for example, the influenza H1 or TTd in this study), and are at very low frequencies in circulation in steady state (24, 25).

Anti-RBD, -S, and -N IgG levels were markedly higher in the convalescent subjects than in non-SARS-CoV-2-exposed subjects, indicating strong induction by SARS-CoV-2 infection. Perhaps notably, the majority of convalescent subjects had higher IgG titers against the S2 than against the RBD. This is particularly surprising because of the accessibility of the RBD to B cells and the expected immunodominance over the S2 subunit (26, 27). Our demonstration of strong anti-S2 IgG production is consistent with the activation of a preexisting population of IgG MBCs against the conserved S2 subunit in the absence of MBCs reactive to the novel RBD. However, we cannot exclude inherent differences in the stability or antigenicity of RBD and S2 reagents as an explanation. In convalescent subjects, IgG levels against the S protein of HCoV OC43 (but not 229E) were significantly higher than in non-SARS-CoV-2-exposed subjects and correlated strongly with anti-S2 IgG levels. These findings support stronger B cell cross-reactivity between the S2 subunits of SARS-Cov-2 and human betacoronaviruses than alphacoronaviruses (8).

Importantly, we demonstrate that SARS-CoV-2 infection generates RBD-reactive and S2-reactive IgG MBCs. Recently, Long et al. (4) found that levels of SARS-CoV-2-reactive Abs, including neutralizing Abs, start to decrease within 8-12 weeks of infection, especially when the infection is asymptomatic. Since MBC populations are maintained for many years, perhaps decades, our findings indicate that MBCs generated by SARS-CoV-2 infection will be available to rapidly generate protective Abs if waning Ab levels allow re-infection to occur (28). Notably, three convalescent subjects in our analysis had undetectable RBD-reactive IgG, but nevertheless had RBD-reactive IgG MBCs. This might reflect MBC production by germinal centers that remained active after recovery from infection (29). The proportion of subjects with MBCs reactive to the HCoVs OC43 and 229E was greater for the convalescent group than the unexposed group, likely reflecting the increase in S2-reactive MBCs in the convalescent group and cross-reactivity with HCoVs. S2-reactive MBC expansion by SARS-CoV-2 infection could enhance protection against a broad range of coronaviruses (23). N-reactive MBC formation in convalescent subjects was less than expected given the large number of subjects with high titers of N-reactive IgG, but additional sampling times are required to confirm this observation.

In conclusion, our analysis investigated Ab and MBC immunity to SARS-CoV-2 in unexposed subjects and individuals soon after recovery from SARS-CoV-2 infection. Findings emphasized the novelty of the SARS-CoV-2 S protein RBD in unexposed subjects. However, IgG reactive to the S2 was widespread in unexposed subjects and likely resulted from exposure to HCoVs. Although our approach was unable to directly identify S2-reactive MBCs in the unexposed subjects, we suggest that these cells are present and strongly contribute S2-reactive IgG early in the response to SARS-CoV-2 infection. The IgG response in SARS-CoV-2 convalescent subjects was also strong against the RBD and, less consistently, against the N protein. Importantly, SARS-CoV-2 convalescent subjects had generated RBD-reactive and S2-reactive IgG MBCs. The RBD-reactive MBCs are likely to provide strong long-term protection if RBD-reactive neutralizing Ab levels wane and re-infection occurs. Additional studies are required to establish the importance of S2-reactive IgG in providing broad anti-coronavirus activity and the influence of expanded S2-reactive MBC populations on a de novo B cell response to the RBD.

## MATERIALS AND METHODS

### Study participants and clinical samples

All study participants were recruited at the University of Rochester Medical Center, Rochester, NY and provided written informed consent prior to inclusion in the studies. The studies were approved by the University of Rochester Human Research Subjects Review Board (protocols 16-0064, 07-0090, and 07-0046) and conducted in accordance with the principles of Good Clinical Practice. A pre-pandemic cohort of 21 healthy donors (median age 48 years, IQR 25-70) were enrolled from 2011-14 (non-SARS-CoV-2-exposed subjects). A cohort of 20 healthcare workers (median age 38 years, IQR 30-52) at Strong Memorial Hospital, Rochester, NY were enrolled in May, 2020. The healthcare workers had not been diagnosed with COVID-19 prior to enrollment. A cohort of 26 non-hospitalized COVID-19 convalescent subjects (9 males and 17 females; median age 49 years, IQR 36-63) was enrolled in May, 2020 and consisted of 22 PCR-confirmed patients and 5 non-PCR-confirmed subjects who were contacts of confirmed cases or displayed COVID-19-like symptoms. The convalescent subjects were sampled 4-9 weeks after symptom onset. Symptoms reported (percent of subjects) were fever (67%) cough (74%), sore throat (48%), stuffy/runny nose (56%), difficulty breathing (52%), fatigue (85%), headache (67%), body aches (67%), nausea/vomiting (19%), and diarrhea/loose stool (41%).

### Recombinant proteins

RBD and stabilized ectodomain S protein from SARS-CoV-2 (isolate Wuhan-Hu-1) were expressed in-house in HEK293 cells using pCAGGS plasmid constructs kindly provided by Florian Krammer (Icahn School of Medicine at Mount Sinai) (7). Baculovirus-expressed S2 subdomain and HEK293 cell-expressed N protein were obtained from Sino Biological (Chesterbrook, PA) and RayBiotech (Peachtree Corners, GA), respectively. Baculovirus-expressed S proteins from seasonal HCoVs OC43 and 229E were obtained from Sino Biological. In-house HEK293 cell-expressed hemagglutinin from egg-derived H1N1 A/California/7/2009 and TTd (MilliporeSigma, Burlington, MA) were used as non-coronavirus control proteins.

### MBC analysis

Measurement of antigen-specific MBCs was essentially performed as described previously (30). Briefly, cryopreserved PBMCs were thawed and rested overnight at 37°C in complete medium. Rested PBMCs were stimulated for 6 days at 1×10^6^ PBMCs/well in 24-well plates to induce MBC expansion and differentiation into ASCs. The stimulation cocktail consisted of complete medium supplemented with 1 μg/ml R848 (Sigma, St. Louis, MO), 10 ng/ml IL-2 (Gibco, Gaithersburg, MD), and 25 ng/ml IL-10 (STEMCELL Technologies, Vancouver, Canada). After stimulation, cells were harvested and pelleted by centrifugation. The undiluted supernatant containing Abs secreted by ASCs generated from stimulated MBC precursors (MPAbs) was collected and stored for analysis by ELISA. Supernatants from unstimulated cultures of rested PBMCs were collected to control for Abs produced by preexisting ASCs. Antigen-specific ASCs in the cell pellet (MASCs) were enumerated by ELISpot assay. For each antigen, 300,000 stimulated PBMCs were analyzed by ELISpot assay and the limit of MASC detection was set at 8 spots (MASCs)/10^6^ PBMCs. Based on ELISpot assay results, antigen-specific MBCs in peripheral blood were quantified as antigen-specific IgG MASCs as a proportion of stimulated PBMCs. Antigen-specific IgG concentrations in MPAb samples (after subtraction of Ab concentrations in supernatants from unstimulated PBMC control cultures) were also used as a measure of the relative size of reactive MBC populations.

### Enzyme-linked immunosorbent assay (ELISA)

Concentrations of Ag-specific serum Abs and MPAbs were measured by ELISA as previously described (30). Briefly, Nunc MaxiSorp 96-well plates (Thermo Fisher, Waltham, MA) were coated overnight with optimized concentrations of antigens. Serially diluted samples were added to blocked plates and incubated for 2 h at room temperature. Alkaline phosphatase conjugated anti-human IgG (clone MT78; Mabtech Stockholm, Sweden) and *p*-nitrophenyl phosphate substrate (Thermo Fisher) were subsequently added to detect bound antigen-specific Abs. Absorbance was read at 405 nm after color development. A weight-based concentration method was used to quantify antigen-specific Ab levels in test samples as described previously (30, 31). Sera from healthy donors and convalescent subjects with high titers for test antigens were used to establish human serum standards. The cutoff for assay positivity was set at approximately 2x the mean OD value for negative wells.

### Statistical analyses

The medians with (q1, q3) were summarized by subject group and compared by the Wilcoxon rank-sum test. Spearman correlation analysis together with corresponding robust regression models was used to assess monotonic associations among Ab responses. Multiple test adjustment was not applied for this explorative study and thus a *P* value < 0.05 was considered significant for all analyses. Statistical analyses were performed using Software SAS 9.4 (SAS Institute Inc, Cary, NC).

## ACKNOWLEDGMENTS

We thank the staff of the University of Rochester Infectious Disease Research Clinic for subject enrollment and sample collection and BEI Resources for providing some of the reagents used in this study.

This project was funded in part with Federal funds from the National Institute of Allergy and Infectious Diseases, National Institutes of Health, Department of Health and Human Services, under CEIRS Contract No. HHSN272201400005C.

The authors declare no conflicts of interest.

